# Denoising diffusion weighted imaging data using convolutional neural networks

**DOI:** 10.1101/2022.01.17.476708

**Authors:** Hu Cheng, Sophia Vinci-Booher, Jian Wang, Bradley Caron, Qiuting Wen, Sharlene Newman, Franco Pestilli

**Affiliations:** Department of Psychological and Brain Sciences, Indiana University, Bloomington, IN 47405, USA; Program of Neuroscience, Indiana University, Bloomington, IN 47405, USA; School of Information Science and Engineering, Shandong Normal University, Jinan, 250358, China; Department of Radiology and Imaging Sciences, Indiana University School of Medicine, Indianapolis, IN 46202; Department of Psychology, University of Alabama, Tuscaloosa, AL 35401; Department of Psychology, The University of Texas at Austin, Austin, TX 78712

**Keywords:** denoising, diffusion weighted imaging, CNN, deep learning

## Abstract

Diffusion weighted imaging (DWI) with multiple, high b-values is critical for extracting tissue microstructure measurements; however, high b-value DWI images contain high noise levels that can overwhelm the signal of interest and bias microstructural measurements. Here, we propose a simple denoising method that can be applied to any dataset, provided a low-noise, single-subject dataset is acquired using the same DWI sequence. The denoising method uses a one-dimensional convolutional neural network (1D-CNN) and deep learning to learn from a low-noise dataset, voxel-by-voxel. The trained model can then be applied to high-noise datasets from other subjects. We validated the 1D-CNN denoising method by first demonstrating that 1D-CNN denoising resulted in DWI images that were more similar to the noise-free ground truth than comparable denoising methods, e.g., MP-PCA, using simulated DWI data. Using the same DWI acquisition but reconstructed with two common reconstruction methods, i.e. SENSE1 and sum-of-square, to generate a pair of low-noise and high-noise datasets, we then demonstrated that 1D-CNN denoising of high-noise DWI data collected from human subjects showed promising results in three domains: DWI images, diffusion metrics, and tractography. In particular, the denoised images were very similar to a low-noise reference image of that subject, more than the similarity between repeated low-noise images (i.e. computational reproducibility). Finally, we demonstrated the use of the 1D-CNN method in two practical examples to reduce noise from parallel imaging and simultaneous multi-slice acquisition. We conclude that the 1D-CNN denoising method is a simple, effective denoising method for DWI images that overcomes some of the limitations of current state-of-the-art denoising methods, such as the need for a large number of subjects for training and accounting for the rectified noise floor.

## 1. Introduction

Neuroanatomical studies rely heavily on diffusion weighted imaging (DWI) sequences because they can provide researchers with estimates of myelinated brain tissue properties *in vivo*. The general approach is to model the diffusion signal and to use parameters of the model fit to infer characteristics about the biological properties of the underlying brain tissue. The earliest and most widely used model is the six-parameter tensor model (i.e. DTI) [1, 2]. A growing body of work is introducing increasingly complex diffusion models with additional parameters that allow researchers to obtain estimates of more nuanced biological properties (e.g., [3–6]). These higher order diffusion models show much promise for advancing our knowledge of the biological properties of brain tissue in living subjects.

Higher order diffusion models are more susceptible to the influence of noise than the standard tensor-based diffusion model because they incorporate DWI images that contain a weak signal. DWI images are collected at a specific diffusion gradient strengths, or *b*-value. The larger the *b*-value, the higher diffusion weighting, the weaker the signal available in the DWI image. The standard tensor-based diffusion model can be fitted using DWI images collected at a *b*-value equal to 1000 s/mm^2^ where the signal is sufficiently strong to overcome several sources of noise. However, the more recently developed higher order diffusion models, such as the models used in Diffusion Kurtosis Imaging (DKI) [7], Mean Apparent Propagator (MAP) MRI [8], and Neurite Orientation Dispersion and Density Index (NODDI) [6], incorporate additional DWI images collected at a higher *b*-value, such as 2500 s/mm^2^. The additional higher *b*-value images contain a weak signal that leads to a more prominent rectified noise floor [9] that, together with the noise present in the image, cause adverse effects on the diffusion metrics derived from these higher order models [10]. Therefore, care must be taken to reduce the effect of noise and the rectified noise floor in DWI images.

Modifications to the diffusion sequence have been proposed to reduce the effects of noise to more effectively capture the weak signal observed at higher *b*-values. One approach is to modify the reconstruction method by adjusting the coil combination scheme. In the case of the multi-channel phased array head coils that are often used in DWI-based neuroanatomical studies, the standard coil combination method is the sum-of-squares (SoS) method [11]. The SoS method weights each channel of the head coil by the pixel value in the channel and can have the unwanted effect of a rectified noise floor when the signal-to-noise ratio (SNR) is small, which becomes especially problematic at high *b*-values when the signal is relatively low. Replacing the SoS coil combination method with a method that incorporates phase information from the coils, such as the adaptive combine (AC) [12] or sensitivity encoding-based (SENSE1) [13] methods, lowers the rectified noise floor, effectively increasing the signal-to-noise ratio (SNR) and facilitating the detection of weak signals. Such modifications are important for recovering the diffusion signal and are particularly important for recovering the diffusion signal at high *b*-values.

The noise floor problem can also be addressed by post-processing methods that are applied after the MRI data have been reconstructed, such as signal transformational frameworks [10, 14, 15] or denoising procedures [14–21]. For instance, the signal-transformational framework reduces the effects of the noise floor by mapping the noisy non-Central Chi signals, such as the Rician distribution often observed in DWI, to noisy Gaussian signals [22]. A critical step of the signal-transformational approach is obtaining the first moment of the non-Central Chi distribution. However, the first moment can only be coarsely estimated through either parametric fitting or spline smoothing of the time course of each voxel. The necessarily coarse estimation of the first moment might introduce bias into the models [10]. An alternative method is to account for the non-Central Chi distribution when fitting diffusion metrics; however, this method can significantly increase the complexity of modeling [23].

The most common post-processing method used to reduce the effects of noise in DWI images is denoising that attempts to reduce the thermal noise in the DWI images. Denoising procedures for DWI images have evolved rapidly in the last decade. The early DWI denoising approach was based on edge-preserved smoothing, such as adaptive smoothing [14, 15] and non-local means [16, 17]. These methods are typically applied on each individual DWI image. Although they are effective at reducing the noise, they tend to blur the images and introduce errors in computing diffusion metrics [19]. The current state-of-the-art methods are based on principal components analysis (PCA) [18, 19] with the assumption of sufficient redundancy of DWI data so that only a number of components can represent the signal-related variations and the rest can be removed as noise. The PCA procedure is often effective in situations where the noise is invariant in the temporal domain, i.e., across DWI volumes, yet fails to effectively address the noise floor effect. For instance, the MP-PCA [19] method fails to reduce the influence of noise amplification in the standard SoS reconstruction [20]. In addition, it is very tricky to set the right threshold to remove the “noisy” component [21]. Although a recent modification of the MP-PCA method resulted in better noise reduction than the original MP-PCA procedure, this recent modification is computationally intensive and not readily implemented on existing data because it is applied to the complex MRI data before reconstruction to avoid complications with the rectified noise floor [24]. Another denoising method, Patch2Self, was proposed to outperform MP-PCA in low-noise datasets by exploiting the statistical independence of noise [25]. However, the simulation result reveals similar poor performance as MP-PCA at the very low SNR (e.g., < 10) typically observed in the high *b*-value DWI images used for higher-order modelling [25]. Therefore, a denoising procedure that can address the rectified noise floor problem in DWI images is needed – one that can be readily implemented on existing data for both high, low, and very low SNR images.

Deep Learning (DL) denoising methods have the potential to overcome some of the limitations of the common denoising technique. DL models have achieved huge success in image denoising [26], as well as other applications, such as speech recognition, object detection, and genomics [27]. Several DL-based methods have been proposed for denoising MRI data, such as convolutional neural networks (CNN) or generative adversarial networks (GAN) [28–32]. One advantage of the DL approach over the PCA approach is that the DL denoising procedure can deal with complex noise behavior [26]. This is particularly important for DWI because DWI datasets contain many images with different image intensities that can have different noise characteristics.

Current DL denoising methods in MRI are mainly applied in the image domain and are aimed at denoising a single image, such as an anatomical image, and, consequently, have been developed in ways that make their application to DWI data impractical. Applying a DL model to DWI data would require a model to be trained for each image in a DWI dataset. Thus, applying the current DL denoising methods to multi-shell DWI data would require a large amount of training data and the training process can be very time consuming. To overcome this problem, recently a method based on a deep image prior was proposed to denoise multiple DWI images simultaneously [33]. While this recently proposed method does reduce the computation time for training, it remains very computationally intensive if the number of DWI images is large (e.g., greater than 30). Given that the multi-shell DWI data often acquired for higher-order models routinely have 60 or more images, this method remains computationally intractable for most researchers.

In this work, we adapted a current DL denoising method for use with DWI data by applying the method to DWI data in the temporal domain, i.e., the time series of the voxels. This approach has two advantages over the more common approach of denoising in the image domain. First, denoising in the temporal domain requires less data than denoising in the image domain because the time course of each voxel is used as a training sample. This makes each voxel a training sample, instead of making each subject a training sample, effectively reducing the amount of required training data to the training required for only one DWI dataset. Second, denoising in the temporal domain is also more likely to retain coherence of the time series for each voxel than denoising in the image domain, in which volumes collected at different timepoints are denoised separately. Retaining coherence of the time series is critical for computing diffusion metrics because the entire time series of a voxel is typically used in diffusion models.

To evaluate this proposed denoising procedure, we first validated the method on simulated DWI data. Then, we evaluated the ability of the method to reduce noise in DWI data after SoS reconstruction, i.e., SoS-related noise. We chose to test our method on SoS-related noise for two reasons: first, current existing denoising methods fail to mitigate the noise floor in high b-value images from SoS reconstruction; second, one can simultaneously reconstruct the image using SENSE1 to serve as the ground truth for training purposes. Finally, we show two possible applications of our method in obtaining low-noise DWI data from deep learning on high-noise DWI data. The first application aims to reduce noise associated with GRAPPA [34]; the second application aims to reduce noise associated with simultaneous multi-slice acquisition (SMS) [35, 36].

## 2. Materials and Methods

### 2.1 The CNN network and Optimization of hyper-parameters

Inspired by the 1D convolutional neural network (CNN) for denoising speeches [37, 38], we constructed a simple 1D-CNN model that has five layers, including two convolutional layers, each followed by a max-pooling layer, and a dense layer (**Figure 1**). The first convolutional layer was given an input of the ‘noisy’ high-noise image and consisted of 16 one-dimensional filtering kernels of size 16. The second convolutional layer consisted of 32 one-dimensional filtering kernels of size 8. The ReLu activation function was used in both convolutional layers. There was 1 max-pooling layer that had a kernel size of 2 with stride 2. In the dense layer, the extracted features of the high-noise image were mapped to the low-noise reference image (**Figure 1**). The model was implemented in Tensorflow v1.14 [39] and python 3.7.3 on a computer with Intel i7-10700 CPU, 32G memory, and Nvidia RTX 2060 GPU.

**Figure 1.**
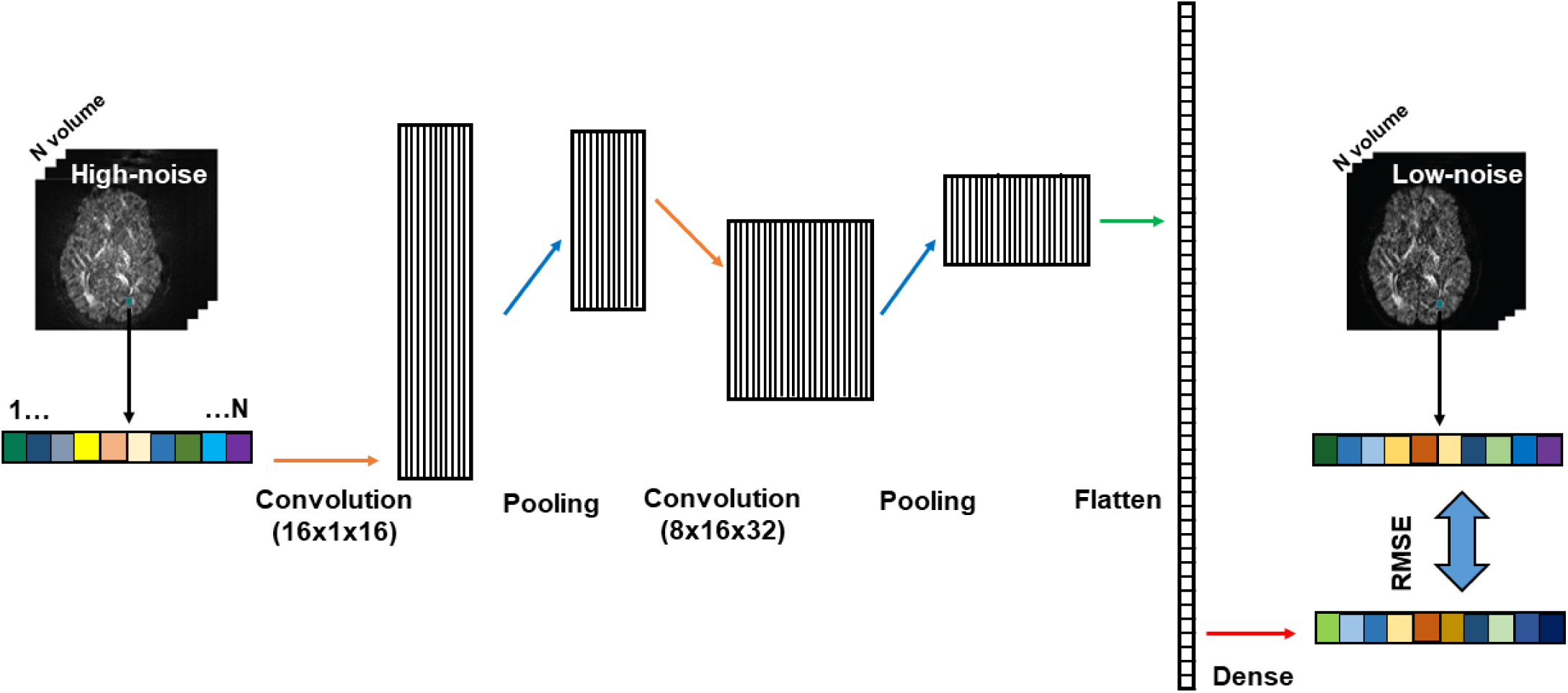
Schematic drawing of the architecture of the 1D CNN model for denoising.

The CNN model uses a stochastic gradient descent algorithm for optimization. This model includes several parameters that, in practice, should be optimized for any specific DWI dataset, namely the number of epochs, learning rate, and batch size. The number of epochs is the number of times that the entire training dataset passes through the model and should be large enough to ensure that the model converges to a point that the error from the model is minimized. The learning rate determines the step size of updating the internal model parameters and needs to be chosen properly to ensure that the training is stable and avoids local minima. The batch size is the number of samples put into the model at one time in the computation that updates the internal model parameters. The updating of the internal model can affect the model error for a specific learning rate.

We selected the number of epochs, learning rate, and batch size empirically based on the training on in vivo data described in **2.2.2**. We kept the number of epochs stable at a high enough value to allow the model to converge to a solution and manipulated two parameters, learning rate and batch size, to find optimal learning parameters. First, we determined the appropriate number of epochs based on a best guess learning rate of 0.001 and batch size of 6000 empirically. We found that the error of the model decreased quickly and stabilized after 6000 epochs for all our data. To be safe, we set the number of epochs to 20000. Second, we determined the optimal learning rate and batch size for the model. We tested learning rate and batch size together because learning rate and batch size often interact in the CNN’s learning procedure. We tested six values for the learning rate [0.00025, 0.0005, 0.001, 0.002, 0.004, 0.008] and four values for the batch size [3000, 6000, 9000, 12000], resulting in 24 tests. For each test, we trained a model on 75% of the voxels from a DWI dataset and, after 20000 epochs, we applied the resulting model to the remaining 25% test voxels. For each combination of learning rate and batch size, the root-mean-square error (RMSE) between the dSoS image and the SENSE1 image in the testing set was used as a measure of performance. We found the optimal learning with the best learning rate of 0.001 and batch size of 6000 (see **Figure S1** in supplementary material).

### 2.2 Validation of Denoising Technique

#### 2.2.1 Validation using simulated data

##### Design

We simulated DWI data to validate the 1D-CNN method for denoising DWI datasets. We simulated a noise-free ground truth and added Gaussian noise to complex MRI data to create both a low-noise and a high-noise dataset to evaluate 1D-CNN denoising for various levels of noise. We then applied both 1D-CNN and MP-PCA [19] denoising to both the low-noise and high-noise datasets to validate the 1D-CNN denoising method against a commonly employed denoising procedure. The design is depicted in **Figure 2A**. We were then able to compare among seven DWI datasets: (1) a noise-free ground truth, (2) low-noise DWI data with SNR = 30, (3) high-noise DWI data with SNR = 10, (4) low-noise after 1D-CNN denoising, (5) high-noise after 1D-CNN denoising, (6) low-noise after MP-PCA denoising, and (7) high-noise after MP-PCA denoising.

**Figure 2.**
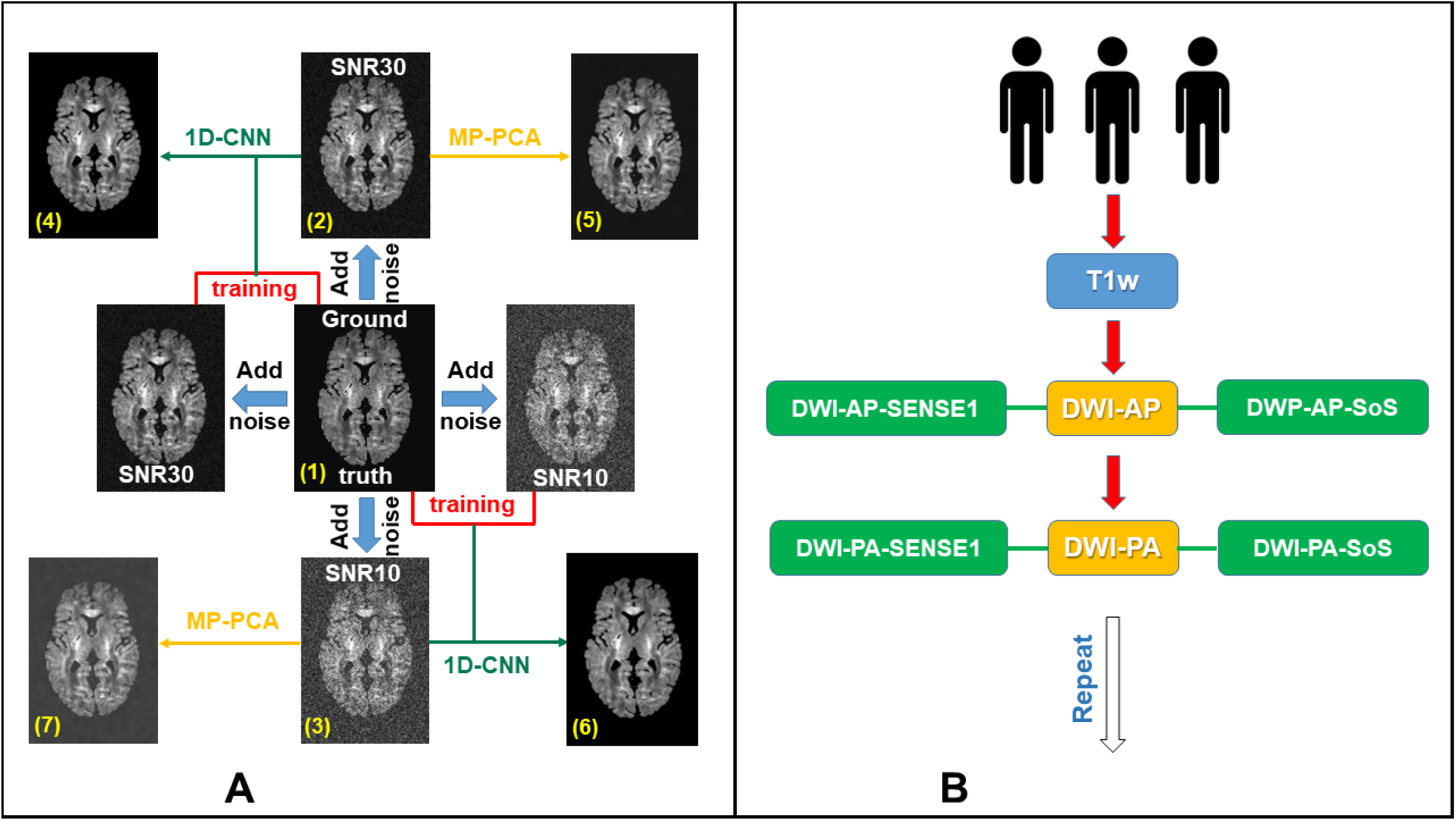
A. We simulated a noise-free ground truth dataset (1) and added noise to create a low-noise dataset with SNR 30 (2) and a high-noise dataset with SNR 10 (3) to evaluate 1D-CNN denoising. The low-noise and high-noise simulated datasets were both denoised using 1D-CNN and MP-PCA, resulting in four additional datasets: low-noise after 1D-CNN denoising (4), low-noise after MP-PCA denoising (5), high-noise after 1D-CNN denoising (6), and high-noise after MP-PCA denoising (7). B. Denoising SoS-related noise using SENSE1 reference data. MRI scans were acquired from 3 subjects for denoising validation. There are two identical runs, each consisting of one T1w scan, two DWI scans with opposite phase encoding directions (DWI-AP and DWI-PA). Each DWI scan is constructed with SoS and SENSE1 coil combines, respectively to generate two sets of images.

##### Data synthesization

We generated five datasets to validate the 1D-CNN procedure for denoising DWI data: (1) a noise-free ground truth, (2) low-noise DWI data with SNR = 30 to be used for model training, (3) high-noise DWI data with SNR = 10 to be used for model training, (4) low-noise DWI data with SNR = 30 to be used for model testing, (5) high-noise DWI data with SNR = 10 to be used for model testing. Four slices of noise-free DWI images were simulated based on the DW-POSSUM framework [40], with 4 volumes at b-value = 0 s/mm^2^ (b0), 90 diffusion directions at b-value = 1000 s/mm^2^ (b1000) and 90 diffusion directions at b-value = 2000 s/mm^2^ (b2000). To generate the low-noise and high-noise DWI datasets, Gaussian noise was added to the real and imaginary channels of the noise-free ground truth using an eight-channel coil model followed by adaptive coil combine. For simplicity, we did not introduce motion, eddy current distortion, or physiological noise into the simulated datasets.

##### Data processing: 1D-CNN

We performed two training procedures, one for the simulated low-noise DWI data and the other for the simulated high-noise DWI data, so that the image pairs used for training the 1D-CNN models were: (1) low-noise DWI data and the noise-free ground truth and (2) high-noise DWI data and the noise-free ground truth. Both 1D-CNN models were trained with the following training parameters: 20000 epochs, learning rate of 0.001, and batch size of 6000. The model trained with the low-noise dataset was applied to the held-out low-noise DWI dataset that was simulated for testing. The model trained with the high-noise DWI dataset was applied to the held-out high-noise DWI dataset that was simulated for model testing.

##### Data processing: MP-PCA

MP-PCA denoising was applied to the simulated low-noise and high-noise DWI data using the “dwidenoise” command in MrTrix3 (https://www.mrtrix.org/). All options were default except for “extent”, whose default setting was changed to 9×9×3 from 9×9×9 to account for our simulated data including only 4 slices.

##### Calculation of diffusion metrics

Diffusion metrics were estimated for all seven comparison images (see ***Validation using simulated data: Design***). Tensor-based diffusion metrics were estimated in FSL (https://fsl.fmrib.ox.ac.uk/fsl/fslwiki/FSL) using the weighted least squares option. NODDI-based diffusion metrics were estimated using the NODDI Matlab toolbox (http://mig.cs.ucl.ac.uk/index.php). We focused on fractional anisotropy (FA), mean diffusivity (MD), intra-cellular volume fraction (ICVF), and orientation dispersion index (ODI) when evaluating the 1D-CNN denoising method.

#### 2.2.2 Validation using *in vivo* data

##### Design

To validate the 1D-CNN denoising method using *in vivo* data, we took advantage of the difference in noise levels that result from different reconstructions of the *k*-space data into image space. More specifically, Sum-of-Squares (SoS) reconstruction of DWI images results in images with higher noise than SENSE1 reconstruction [13] and, crucially, the high-noise and low-noise pair are derived from the exact same image.

Three human participants were recruited. We collected two DWI images for each subject. The first image was reconstructed using both the sum-of-squares (SoS) and also the SENSE1 coil combination methods, resulting in an SoS and SENSE1 reconstruction of the same image. The second image was reconstructed using only the SENSE1 coil combination method and was used as a test-retest repeat of the SENSE1 reconstruction of the first image. We then applied 1D-CNN denoising to the SoS image to generate a denoised SoS image (dSoS). This procedure resulted in four comparison datasets: (1) SoS, (2) SENSE1, (3) dSoS, and (4) SENSE1-repeat.

We took several steps to test the effects of 1D-CNN denoising on diffusion metrics in *in vivo* data. First, we computed FA and MD from the tensor model using all DWI images. Second, we computed OD and ICVF from the NODDI model to evaluate the effects of the denoising procedure on higher-order diffusion models. Third, we estimated each metric, FA, MD, OD, and ICVF, in the gray matter and white matter separately. As the DWI signal attenuation from diffusion weighting varies with tissue types, which ultimately affects the SNR, it is worth investigating if the effectiveness of 1D-CNN denoising procedure was dependent on the tissue type. Finally, we tested the performance of 1D-CNN by visually inspecting the images of each diffusion metric calculated from SoS, dSoS, and SENSE1 data, plotting histograms of each metric calculated from SoS, dSoS, and SENSE1 data, and computing correlations between the diffusion metrics calculated from the SENSE1 data and the SoS and dSoS data. Given that SoS-related noise was expected to be particularly problematic at higher *b*-values, we expected the denoising procedure to impact FA/MD and ICVF estimates derived from multi-shell data that contains high *b*-value images. We expected the histogram and correlation analyses to demonstrate that the proposed denoising procedure results in diffusion metrics that are more similar to the metrics computed on the SENSE1 standard than metrics computed on the original SoS data and, further, than metrics computed on a SENSE1 repeat.

##### Data acquisition

We collected DWI data from three subjects using the Human Connectome Project (HCP) Lifespan protocol [41] on a Siemens 3T Prisma (Siemens Healthineers, Erlangen, Germany). The procedure began with one anatomical T1w image followed by a series of DWI images and a repeat of the T1w and diffusion images. See **Figure 2** for details of the DWI data acquisition procedure. T1w images were acquired using the 3D MP-RAGE pulse sequence: TR/TE=2400/2.3 ms, TI=1060 ms, flip angle=8, matrix=320×320, bandwidth=210 Hz/pixel, iPAT=2, resulting in 0.8 mm isotropic resolution. DWI images were collected in both AP and PA phase encoding directions with the following parameters: TR/TE = 3470/87 ms, 72 slices, 1.5 mm isotropic resolution, 37 gradient directions with b-value = 1000, 2500 s/mm^2^ plus 6 b0 images, resulting in a total of 80 images, SMS acceleration factor = 4. In addition, we collected one repetition of the AP and PA DWI images for each subject. Crucially, each DWI images was reconstructed with both SoS and SENSE1 [13] coil combination methods so that the SoS and SENSE1 data used for the model training and testing (described below) originated from the same k-space data.

##### Anatomical image processing

We applied standard preprocessing steps to the T1w images. Anatomical images were aligned to the AC-PC plane with an affine transformation using HCP preprocessing pipeline [42] as implemented in the HCP AC-PC Alignment App on brainlife.io (bl.app.99) to the standard MNI152 adult template. AC-PC aligned images were then segmented using the Freesurfer 6.0 [43] as implemented in the Freesurfer App on brainlife.io (bl.app.0) to generate the cortical volume maps with labeled cortical regions according to the Destrieux 2009 atlas [44]. We collected on an additional repeat T1w image for each subject.

##### Diffusion image processing

Processing of the DWI data occurred in three steps before the data were used to calculate various diffusion metrics and run fiber tracking. First, we applied minimal preprocessing to the DWI images to prepare them for the training and optimization procedure. AP phase-encoded and PA phase-encoded images were combined using FSL Topup & Eddy to correct for susceptibility distortion correction and inter-volume subject motion (brainlife.app.287). This step combined AP and PA images, resulting in one SoS image, one SENSE1 image, and one SENSE1 repeat for each subject. A mask of the brain was extracted from the DWI with *b*-values equal to 0 s/mm^2^. All following analyses were conducted using only voxels within this brain mask.

Second, we trained the 1D-CNN model using the SoS and SENSE1 data of each subject with the optimal parameters (20,000 epochs, learning rate = 0.001, batch size = 6000), resulting in three sets of model coefficients, one for each subject. We then applied the trained model obtained for each subject to the SoS data of the other two subjects separately in a round-robin style, resulting in two sets of dSoS data for each subject. With the addition of the dSoS data, data for each subject included one SoS image, one SENSE1 image, one SENSE1 repeat, one dSoS image that was denoised using the model coefficients of a second subject, and one dSoS image that was denoised using the model coefficients of the remaining subject.

Third, additional preprocessing steps were applied to each image separately. Additional preprocessing steps included PCA denoising, Gibbs deringing, bias field correction, and Rician noise removal available in the Mrtrix3 preprocessing pipeline [45] as implemented in the Mrtrix3 Preprocess App on brainlife.io (bl.app.68). PCA denoising and Gibbs deringing procedures were performed first. The preprocessed DWI data and gradients were then aligned to each subject’s ACPC-aligned anatomical image using boundary-based registration (BBR) in FSL [46].

##### Calculation of diffusion metrics

To evaluate the effect of denoising on tensor-based diffusion metrics and diffusion metric derived from higher order diffusion models, we fit both the tensor model [2] and the Neurite Orientation Dispersion and Density Index (NODDI) model [6] to the processed DWI data. Fractional anisotropy (FA) and mean diffusivity (MD) were estimated from the tensor model in FSL using the weighted least square option. We measured intra-cellular volume fraction (ICVF) by fitting the NODDI model on multi-shell data using the NODDI Matlab toolbox developed by UCL Microstructure Imaging Group (http://mig.cs.ucl.ac.uk/index.php).

##### Tractography

To evaluate the effect of denoising on tractography, tractography was performed on each DWI dataset to generate streamlines. We used constrained spherical deconvolution (CSD) to model the diffusion tensor for tracking [47, 48]. Tracking with the CSD model fit was performed probabilistically using the tractography procedures provided by Mrtrix3 Anatomically constrained Tractography (ACT) [49, 50] implemented in brainlife.io (bl.app.101). We generated 150000 streamlines at L_max_ = 8 and maximum curvatures = 35 degrees. The tractogram was then segmented into major white matter tracts using a recently developed, automated segmentation approach [51] implemented in brainlife.io (brainlife.app.188). Streamlines that were shorter than 10 mm or longer than 200 mm were excluded. Streamlines that were more than 4 standard deviations away from the centroid of each tract and/or 4 standard deviations away from the tract’s average streamline length were considered aberrant streamlines and were removed using the Remove Tract Outliers App on brainlife.io (brainlife.app.195). Streamline counts were obtained using the Tract Statistics App on brainlife.io (brainlife.app.189). See **Table 1** for more information on the Brainlife applications used.

**Table 1.** Data, description of analyses, and web-links to the open source code and open cloud services used in the creation of this dataset can be viewed in their entirety here: https://doi.org/10.25663/brainlife.pub.25.

| APPLICATION | GITHUB REPOSITORY | OPEN SERVICE DOI | GIT BRANCH |
| --- | --- | --- | --- |
| HCP AC-PC ALIGNMENT | https://github.com/brain-life/app-hcp-acpc-alignment | doi.org/10.25663/bl.app.99 | 1.4 |
| FREESURFER SEGMENTATION | https://github.com/brainlife/app-freesurfer | doi.org/10.25663/bl.app.0 | 1.7 |
| DISTORTION AND MOTION CORRECTION | https://brainlife.io/app/5e6e72838a20890d8a8e96af | doi.org/10.25663/brainlife.app.287 | master |
| DMRI PREPROCESSING | https://github.com/brain-life/app-mrtrix3-preproc | doi.org/10.25663/bl.app.68 | 1.5 |
| TRACTOGRAPHY | https://github.com/brain-life/app-mrtrix3-act | doi.org/10.25663/bl.app.101 | 1.3 |
| TRACT SEGMENTATION | https://github.com/brainlife/app-wmaSeg | doi.org/10.25663/brainlife.app.188 | 3.3 |
| TRACT CLEANING | https://github.com/brainlife/app-removeTractOutliers | doi.org/10.25663/brainlife.app.195 | 1.3 |
| TRACT STATISTICS | https://github.com/brainlife/app-tractographyQualityCheck | doi.org/10.25663/brainlife.app.189 | 1.2 |

To evaluate the effect of 1D-CNN denoising on tractography, we counted the number of streamlines in several major white matter tracts and calculated the difference in the streamline counts for these tracts between the gold standard SENSE1 reconstruction and the other DWI images using the multi-shell data: SoS, dSoS, and the SENSE1 repeat. Because the noise in DWI data blurs the orientation dispersion function (ODF), resulting in lower FA values and less accurate tractography, we expected that tractography would perform best in the SENSE1 images followed by the dSoS images and then the original SoS image. We, therefore, expected the SENSE1-dSoS difference would be smaller than the SENSE1-SoS difference, suggesting that 1D-CNN denoising positively affected tractography.

### 2.3 Application

The high-noise and low-noise dataset pairs used in the simulation and *in vivo* studies described above came from the same base image; however, it is likely in human subject research that the high-noise and low-noise dataset pair will come from different base images. To assess the feasibility of the 1D-CNN method for denoising high-noise DWI data using a low-noise dataset derived from a different base image, we conducted two experiments. In the first experiment, we denoised a high-noise DWI dataset acquired with GRAPPA [34] by training the 1D-CNN model on a GRAPPA (i.e., high-noise) and a non-GRAPPA (i.e., low-noise) pair. In the second experiment, the high-noise dataset was an simultaneous multi-slice (SMS) [35, 36] dataset while the low-noise dataset was without SMS. GRAPPA and SMS were selected because they are normally used in DWI to decrease acquisition time. The detailed acquisition parameters are shown below.

#### Experiment 1, GRAPPA

Three subjects were scanned on a Prisma scanner for two consecutive runs with the following parameters: TR/TE = 5400/88.2 ms, 72 slices, 1.5 mm isotropic resolution, 37 gradient directions with b-value = 1000, 2500 s/mm^2^ plus 6 b0 images, simultaneous multi-slice (SMS) acceleration factor = 2, resulting a total of 80 images. GRAPPA with acceleration factor 2 (iPAT2) was used in the first run but not in the second run. Therefore, the SNR of the images in the first run with iPAT equal to 2 is lower than that in the second run with iPAT equal to 1.

#### Experiment 2, SMS

Three subjects were scanned on a Prisma scanner for two consecutive runs with the following parameters: TR/TE = 3470/87 ms, 1.5 mm isotropic resolution, 37 gradient directions with b-value = 1000, 2500 s/mm^2^ plus 6 b0 images, resulting a total of 80 images. The SMS option with an acceleration factor of 4 was used in the first run, resulting in 72 slices, but not in the second run, resulting in 18 slices.

#### 2.3.3 Data processing

All images were processed in FSL first for Eddy current/motion correction. For Experiment 1, the susceptibility-induced distortion was slightly different between the two runs. For Experiment 2, there was also a slight anatomical mismatch between the two runs. Therefore, a nonlinear transformation was performed on the low-noise images to align them with the high-noise ones. The training of 1D-CNN for denoising was between the low-noise and high-noise datasets of subject 2 and the model was applied to the other two subjects. Again, we chose learning rate of 0.001 and batch size of 6000. The fitting converged quickly before reaching 20000 epochs.

## 3. Results

### 3.1 Validation of 1D-CNN on simulated data

**Figure 3** shows the difference between denoised image and the noisy image as well as the difference between denoised image and the ground truth of a representative slice with b-value 2000 s/mm^2^ from the SNR=10 dataset. MP-PCA removed some noise but was unable to restore the signal to the ground truth, as manifested in the CSF region. In contrast, 1D-CNN mapped the noisy image to the ground truth, although the removed noise shows some structures of the brain. **Figure 4** compares the resultant diffusion metrics of FA, MD, and ICVF for noisy images with SNR 10 and 30, the corresponding denoised images using MP-PCA and 1D-CNN, and the ground truth. The noise floor led to an underestimation of FA and MD and an overestimation of ICVF that becomes more prominent as SNR decreases. MP-PCA denoising alleviated that effect on FA and MD but not on ICVF. In contrast, 1D-CNN significantly improved the accuracy of MD and ICVF. At SNR = 30, however, there is little difference between the results of MP-PCA, 1D-CNN, and uncorrected.

**Figure 3.**
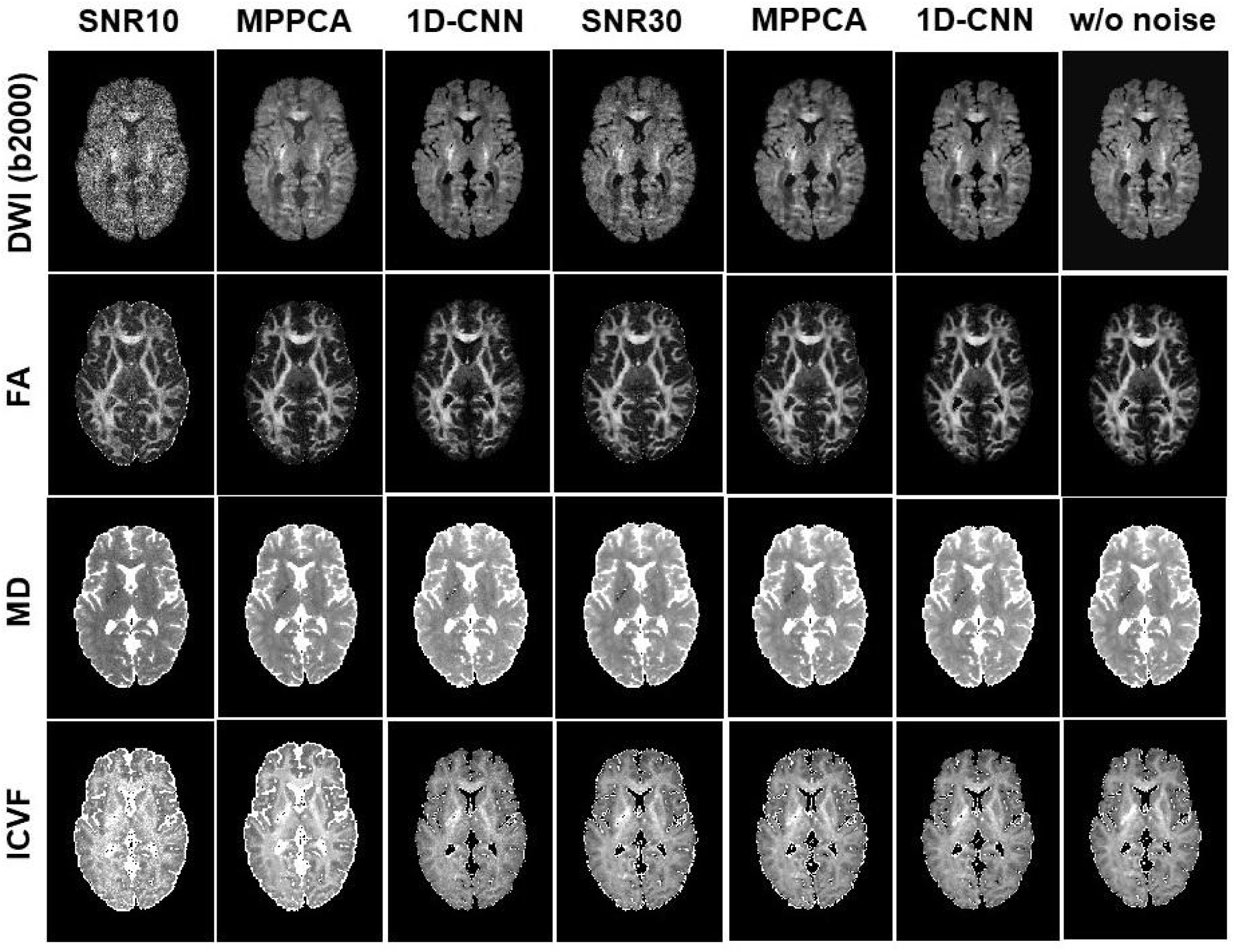
The noisy DWI images (SNR 10 and 30), corresponding denoised images with MP-PCA and 1D-CNN, and the ground truth, along with derived FA, MD, and ICVF maps of a representative slice from simulation.

**Figure 4.**
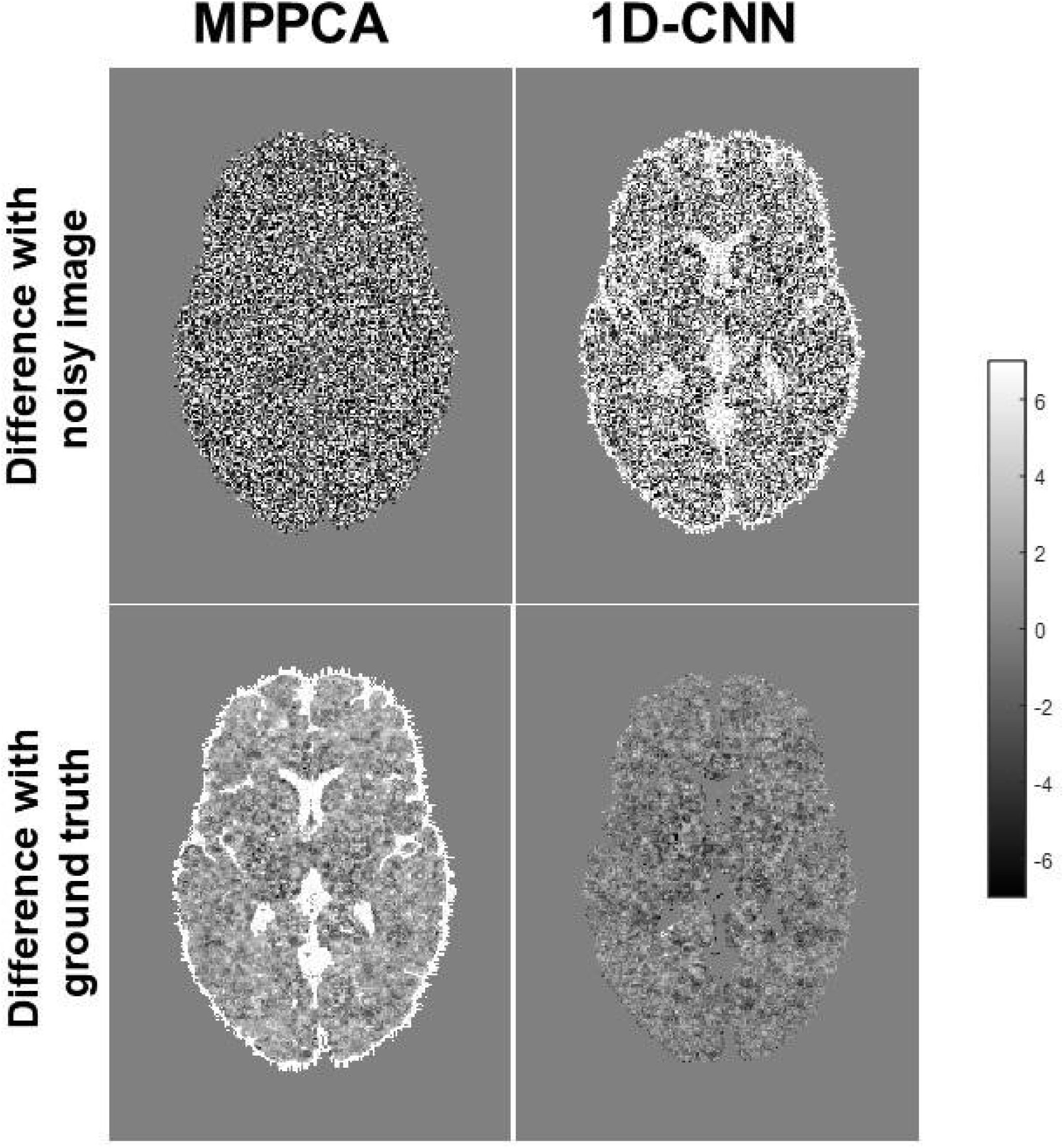
Difference (arbitrary unit) between denoised image and original noisy image (top) and the ground truth (bottom) for MP-PCA and 1D-CNN from simulation.

### 3.2 Evaluation of 1D-CNN on *in vivo* data

The performance of 1D-CNN was evaluated in the *in vivo* dataset in three domains: DWI images, diffusion metrics, and tractography.

In the DWI image domain, the denoised SoS (dSoS) images were compared to the SENSE1 gold standard through visual inspection and also by evaluating the correlation between the DWI signal in dSoS and SENSE1 images. Visual inspection of the DWI images indicated that 1D-CNN resulted in DWI images that were more similar to the gold standard SENSE1 images than the original SoS reconstructed images. The denoised SoS (dSoS) images appeared less noisy than the original SoS images. A representative slice of the SoS image, the SENSE1 image, and the dSoS image for one subject are shown in **Figure 5a** for *b* = 1000 s/mm^2^ and *b* = 2500 s/mm^2^. The average correlation coefficients between SENSE1 and dSoS images were computed for each *b*-value (**Figure 5b**). The correlation between SENSE1 and SoS images was improved after denoising in both low and high b-value images, with high b-value images improved the most. Notably, the correlations were much greater than those between SENSE1 repeat scans (*r* = 0.6 – 0.8).

**Figure 5.**
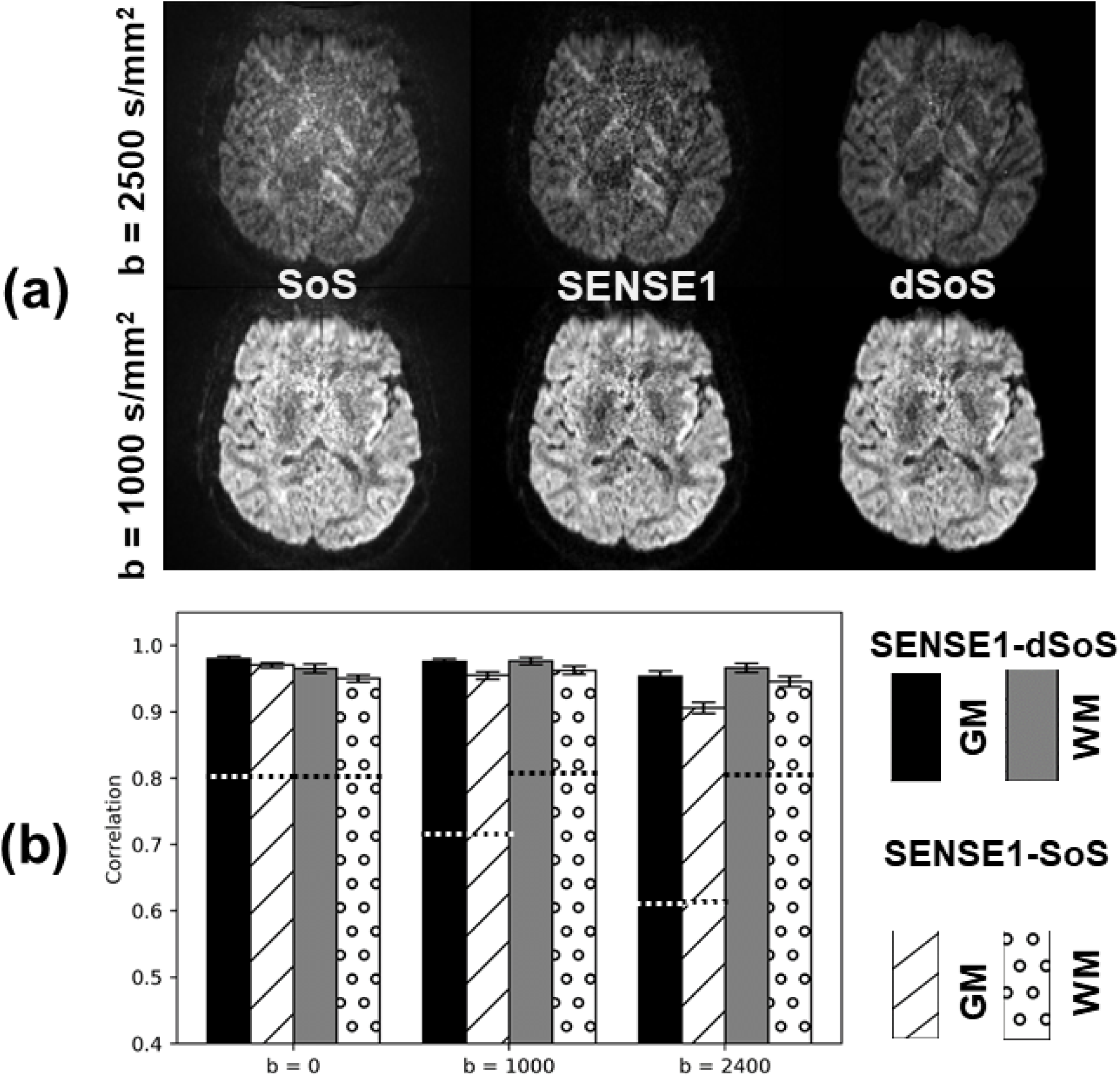
(a) A representative slice of the SoS image, the SENSE1 image, and the denoised SoS (dSoS) image for b = 1000 s/mm^2^ and b = 2500 s/mm^2^; (b) The average correlation coefficients between SENSE1 and dSoS images, and between SENSE1 and SoS images, for gray matter (GM) and white matter (WM). The dashed lines indicate the magnitude of correlation coefficients between two repeated scans with SENSE1 (SENSE1 inter-scan) as a reference.

In the diffusion metrics domain, there was little difference for FA among the SoS, SENSE1, and dSoS data based on visual inspection of the maps and histograms, as shown in **Figure 6**. However, MD, OD and ICVF were positively affected by the denoising procedure. The histograms revealed that OD and ICVF calculated from the SoS data were overestimated while MD was underestimated relative to the SENSE1 standard, and that effect was mitigated substantially by denoising (**Figure 6**). Comparing the correlations among SoS, SENSE1, and dSoS images revealed that the SENSE1 data were more highly correlated with the dSoS data (OD: 0.9744 in GM and 0.9790 in WM, ICVF: 0.9592 in GM and 0.9180 in WM) than with the SoS data (OD: 0.9558 in GM and 0.9615 in WM, ICVF: 0.8529 in GM and 0.8499 in WM) (**Table 2**).

**Figure 6.**
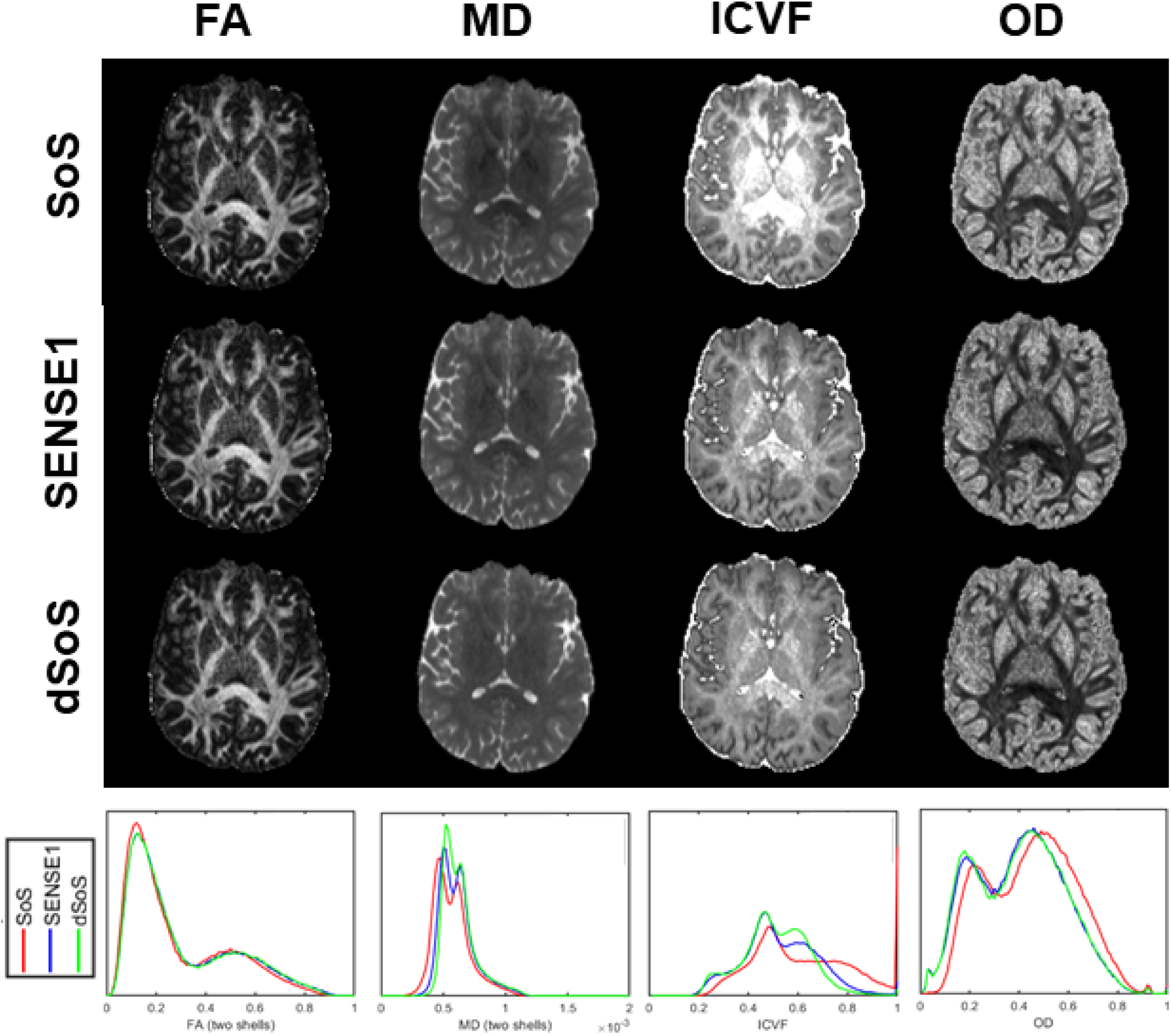
Different diffusion metrics (FA, MD, ICVF, and OD) computed from SoS, SENSE1, and dSoS images showing on a representative slice. The corresponding histograms are shown in the bottom. For ICVF and OD, the dSoS data (green) is more similar to the gold standard SENSE1 data (blue) than the original SoS data (red). The benefit of our denoising procedure is most evident in ICVF and OD, the two measures that are derived from higher-order diffusion models that rely on high *b*-value images, although some improvement can also be seen in the MD value.

**Table 2.** Correlations of the tensor-based diffusion metrics (i.e., FA, MD) and NODDI-based diffusion metrics (i.e., OD, ICVF) between SENSE1-SoS, SENSE1-dSoS, and SENSE1-SENSE1 repeat.

|  | **SENSE1-SOS** | **SENSE1-DSOS** | **SENSE1 –**<br>**REPEAT** |
| --- | --- | --- | --- |
| **FA (GM)** | 0.949 | 0.961 | 0.924 |
| **FA (WM)** | 0.957 | 0.964 | 0.941 |
| **MD (GM)** | 0.817 | 0.920 | 0.828 |
| **MD (WM)** | 0.572 | 0.808 | 0.719 |
| **OD (GM)** | 0.956 | 0.974 | 0.867 |
| **OD (WM)** | 0.962 | 0.979 | 0.932 |
| **ICVF (GM)** | 0.853 | 0.959 | 0.881 |
| **ICVF (WM)** | 0.850 | 0.918 | 0.850 |

In the tractography domain, we counted the number of streamlines in several major white matter tracts and calculated the difference in the streamline counts for these tracts between the gold standard SENSE1 reconstruction and the other DWI images using the multi-shell data: SoS, dSoS, and the SENSE1 repeat. These differences are shown in **Figure 7** for nine major white matter tracts: IFOF (left and right), ILF (left and right), Forceps Major, Arc (left and right), and SLF 1 and 2 (left and right). The SENSE1-dSoS differences were smaller or no different than the SENSE1-SoS differences for all tracts and, importantly, the SENSE1-dSoS differences were never greater than the SENSE1-SoS differences, indicating that the tractography algorithm did not perform worse on the dSoS data than on the original SoS data.

**Figure 7.**
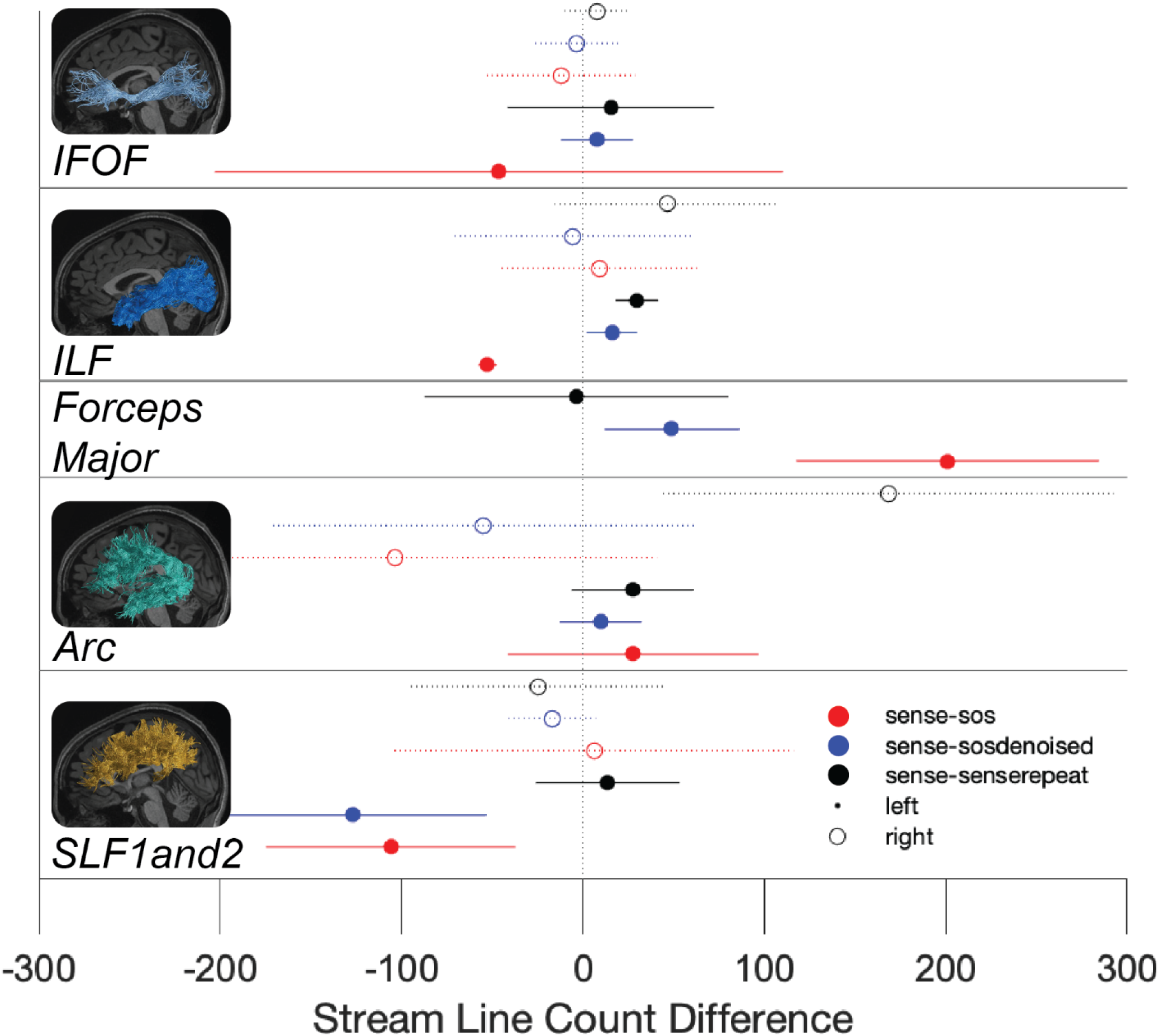
The difference in the number of tractography streamlines between SENSE1 and SoS, SENSE1 and dSoS, and SENSE1 repetitions for nine bundles: IFOF (left and right), ILF (left and right), Forceps Major, Arc (left and right), and SLF 1 and 2 (left and right). The x-axis is the average difference in streamline count between the comparison (i.e., SENSE1-SoS (red), SENSE1-dSoS (blue), SENSE1-SENSE1 repeat (black)). The error bars are standard deviation.

### 3.3 Effect of training samples

The proposed denoising procedure requires only data from one subject for training. This is a strength of the procedure because it allows researchers to perform 1D-CNN denoising without having to collect an extremely large amount of data. However, training the model on only one subject presents the potential concern that the model coefficients might receive some bias related to the training subject’s data such that the application of the trained model to another subject’s data would effectively bias the second subject’s data towards the training subject. This would be problematic for the majority of neuroanatomical studies because such studies often aim to interrogate individual or group differences. If 1D-CNN biases the denoised data towards a single subject, i.e., the training subject, then any real differences present among subjects or between groups would be more difficult to detect.

The impact of this potential bias of the training subject on the denoised image was interrogated by computing the correlation coefficients between diffusion metrics of dSoS images denoised using models trained from different subjects’ data. If the dSoS images are influenced by the training subject, then we would expect to find a low correlation if the dSoS images were influenced by the spatial characteristics of the training data because the training data from different subjects are totally independent.

**Table 3** lists the correlation coefficients of three diffusion metrics (FA, ICVF, OD) derived from denoised SoS data of each subject using the other two subjects as training data, respectively. For FA and OD, the correlation coefficients in GM and WM were all greater than 0.99. For MD, the correlation coefficients were all greater than 0.99 in GM and 0.96 in WM. The correlation coefficients were slightly lower for ICVF, but the values (> 0.96 in GM and > 0.94 in WM) are still much higher than those between two repeated scans. The corresponding scatter plots of these four diffusion metrics are shown in **Figure S2** in the **Supplementary Material**. Although the diffusion measures differ with different training dataset, the high correlation coefficients suggest that the choice of subject used as a training dataset has a negligible impact on denoised images.

**Table 3.** SENSE1-dSoS correlations of diffusion metrics (FA, MD, OD, and ICVF). For the dSoS data used in this table, the model was trained on data that was merged from the other two subjects. The dSoS data for SUBJ 1, for example, was obtained by applying a model trained on the combined data of SUBJ 2 and SUBJ 3 to SUBJ 1’s SoS data.

|  | FA |  | MD |  | OD |  | ICVF |  |
| --- | --- | --- | --- | --- | --- | --- | --- | --- |
|  | GM | WM | GM | WM | GM | WM | GM | WM |
| **SUBJ 1** | 0.998 | 0.997 | 0.995 | 0.975 | 0.994 | 0.995 | 0.978 | 0.943 |
| **SUBJ 2** | 0.990 | 0.997 | 0.993 | 0.978 | 0.991 | 0.994 | 0.981 | 0.962 |
| **SUBJ 3** | 0.997 | 0.997 | 0.991 | 0.966 | 0.991 | 0.993 | 0.972 | 0.941 |

### 3.4 Applications

As denoising validation from simulated data and in vivo data indicated that denoising has little impact on FA and OD, we only show the effect on MD and ICVF in the two application experiments. **Figure 8** shows the original high-noise image, low-noise image, and denoised image using 1D-CNN along with derived MD and ICVF maps of a representative slice in Experiment 1 (iPAT2 vs. iPAT1) for b-value 2500 s/mm^2^. Similarly, an underestimation of MD and a ceiling effect of ICVF were observed for high-noise data were mostly corrected after 1D-CNN denoising. **Figure 9** shows the original high-noise image, low-noise image, and denoised image using 1D-CNN along with the derived MD and ICVF maps of a representative slice in Experiment 2 (SMS4 vs. SMS1) for b-value 2500 s/mm^2^. Again, an underestimation of MD and a ceiling effect of ICVF were observed for high-noise data that were corrected after 1D-CNN denoising. 1D-CNN denoising not only revealed more details in the low *b*-value image that were present in the high *b*-value images but also made the MD and ICVF values closer to those derived from the low-noise dataset.

**Figure 8.**
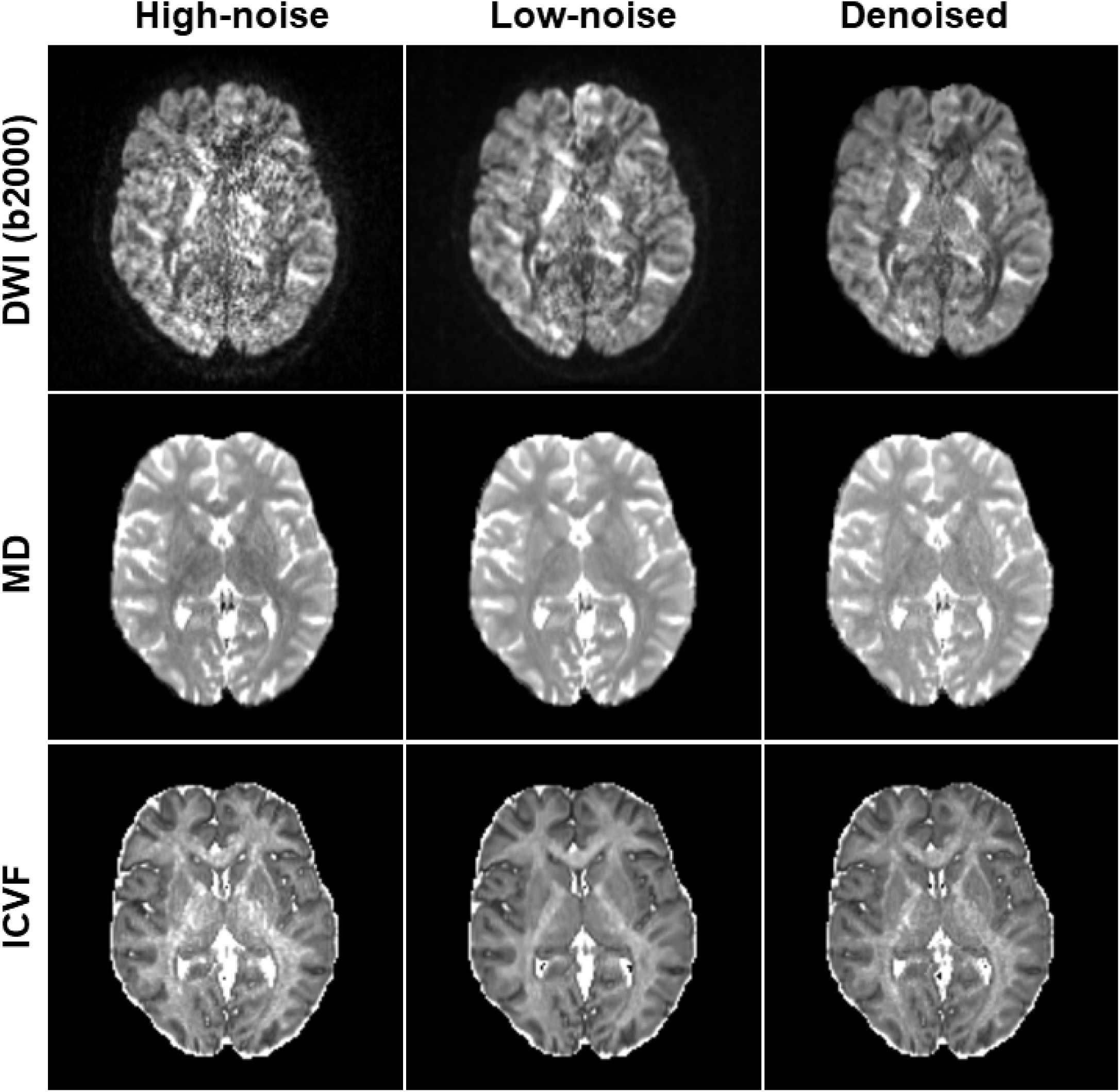
The original high-noise image, low-noise image, and denoised image using 1D-CNN along with derived MD and ICVF maps of a representative slice in Experiment 1 (iPAT2 vs. iPAT1) for b-value 2500 s/mm^2^.

**Figure 9.**
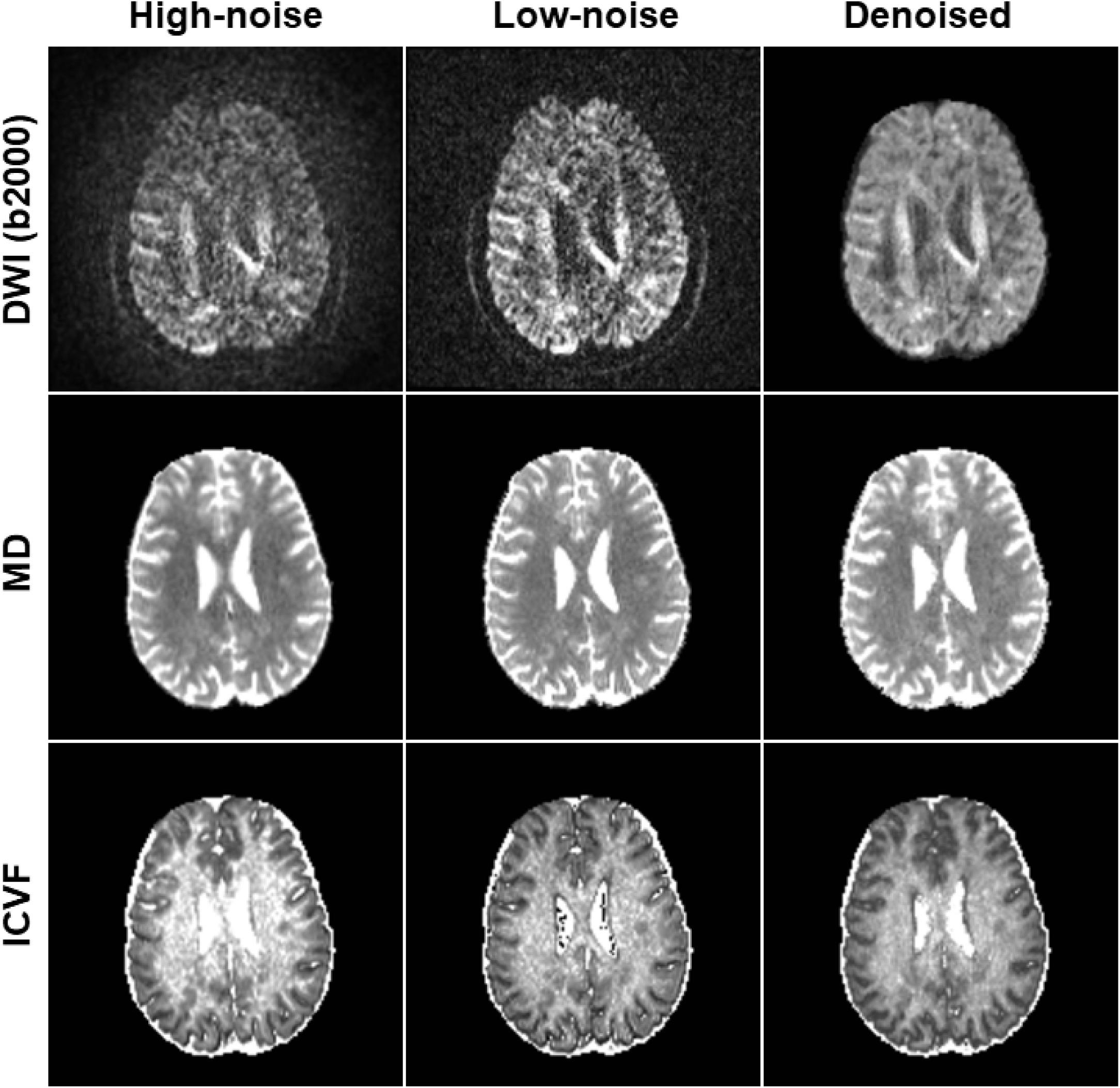
The original high-noise image, low-noise image, and denoised image using 1D-CNN along with derived MD and ICVF maps of a representative slice in Experiment 2 (SMS4 vs. SMS1) for b-value 2500 s/mm^2^.

## 4. Discussion

The behavior of noise in DWI images varies across images. For those with high SNR the noise is exhibited as Gaussian noise; for those with very low SNR (say, < 10), rectified noise floor can appear. Rectified noise floor can severely lower the accuracy in estimating diffusion metrics sensitive to diffusion properties at high b-values, such as ICVF from NODDI, mean kurtosis from DKI, and various measures from MAP-MRI. Most DWI denoising methods work well in removing the Gaussian noise but not the rectified noise. Our primary goal was to develop a practical method for removing noise in DWI data that is not removed adequately by currently available denoising methods. We were particularly interested in developing a method that would be effective for multi-shell DWI data that contain high *b*-values that are required for higher order diffusion models. Using simulated data and SoS noise in *in vivo* data as examples, we demonstrated that a deep learning method was able to correct for rectified noise floor in DWI images that cannot be corrected using state-of-the-art DWI denoising methods. Our method was based on a one-dimensional CNN deep-learning network applied to the time course of DWI data, substantially reducing the amount of training data required for denoising relative to current DL denoising procedures. With only one dataset as the training sample (i.e., one subject’s worth of DWI data), the 1D-CNN method effectively removes noise for any dataset acquired the same way as the training data.

The proposed 1D-CNN denoising method is essentially a regression model that fits the low-noise images from high-noise images rather than fitting the noise. Therefore, it can deal with more complicated noise behavior. The simulation results demonstrated that the ultimate target of restoring a noisy image to its noise-free reference image can be accomplished with 1D-CNN, a feature that sets it apart from other state-of-the-art denoising methods (e.g., MP-PCA). In denoising SoS-related noise using SENSE1 data as the ground truth, 1D-CNN reduced the effects of noise so that the denoised DWI images and subsequently derived diffusion metrics were more similar to the ground truth than a repeated scan. The voxel-wise correlations of the image intensity between denoised SoS data and SENSE1 were above 0.95 for all subjects tested, much higher than the correlations between repeated scans using the SENSE1 coil combine mode (**Figure 5**). Similarly, the voxel-wise correlations between the diffusion metrics computed from dSoS and SENSE1 data were also very high and always greater than the voxel-wise correlations between the SENSE1 and SENSE1-repeat scans (**Table 2**). Both results indicate that the proposed denoising method can substantially reduce SoS-related noise to a level where the diffusion signal can be recovered and modelled with as much fidelity as if the DWI data were originally collected with a SENSE1 coil combine method.

We did not observe substantial benefit of 1D-CNN denoising for FA. This is probably because FA is more determined by the directionality rather than diffusion signal at high b-value. The noise floor is particularly problematic for higher order diffusion models that rely on high *b*-value images. The correlation coefficients of all diffusion metrics were either marginally or significantly improved after denoising – an effect that was most evident for measurements obtained using higher order diffusion models (**Figures 4**, **7** and **Table 2**).

Tractography results were generally in line with our expectations that the denoising procedure would improve tractography performance. The difference between streamline counts for the SENSE1 standard and the dSoS tractogram was smaller or no different than the difference between the SENSE1 standard and the SoS tractrogram for each tract (**Figure 7**). However, these results also demonstrated that SoS-related noise is not likely problematic for tractography performance because the difference in streamline counts between the SENSE1 standard, the SoS tractogram, and the dSoS tractogram were very small relative to the total number of streamlines (i.e., < 1%). We had expected tractography performance to be negatively affected by the SoS-related noise; however, little to no effect in tractography performance is consistent with our results that demonstrated little to no effect of SoS-related noise on FA, which is closely related to the directionality of diffusion.

### Effect of training data

In theory, the training data must contain all the characteristic time courses of the DWI data to be denoised, which are related to the tissue microstructure and coil configuration. This requires a vast amount of training data. One of the advantages of the proposed method is that it only needs a few DWI datasets because each DWI dataset consists of many training samples (∼5×10^5^ for HCP protocol). Although it is possible to use multiple datasets to train the model, we have shown that even one DWI dataset acquired with the same protocol is adequate for the training purpose. In addition, although different training data leads to deviated denoising results, as illustrated by the scatter plots of diffusion metrics in **Figure S2**, the choice of training data does not affect the results dramatically, as suggested by the high correlation between resultant diffusion metrics using different training data for denoising (**Table 3**). Therefore, the training can be carried out using one representative dataset in practice. However, the hyper-parameters of the model need to be tuned separately for each protocol because the choice of learning rate and batch size affects the performance of the algorithm using gradient descent for optimization. We expect different values for the optimal learning rate and batch size for different acquisition protocols. It seems that for an optimal learning rate, there is some freedom in setting the batch size (e.g., learning rate 0.01 and 0.02 in **Figure S1**).

### Possible applications

The python script for denoising can be downloaded from https://github.com/huchengMRI/DWI-SoS-denoising. We provided codes for both Tensorflow 1.1 and 2.0. There could be many applications of 1D-CNN in denoising DWI data. The first one is retrospectively correcting DWI data with high *b*-values that was reconstructed with SoS coil combine. The researcher must collect at least one DWI dataset with the same acquisition scheme as the data-to-be-denoised. The images must then be reconstructed using, one, the Sum-of-squares (SoS) reconstruction method and, two, the SENSE1 reconstruction method. These are the datasets to be fed into the CNN architecture. The datasets can be divided into training and validation parts to find the optimal hyper-parameters (e.g., learning rate, batch size, number of epochs) for the specific DWI acquisition scheme. Then, the trained model can be applied to any high-noise dataset for denoising as long as it was acquired with the same acquisition parameters.

Parallel imaging techniques have been routinely used in DWI scans to save time, reduce geometric distortion, and improve spatial resolution despite an SNR penalty [52]. The SNR penalty is proportional to the square root of the acceleration factor multiplied by a g-factor that is mainly determined by the coil configuration and acceleration factor [52]. Experiment 1 suggested practical applications of using 1D-CNN to reduce the effect of noise floor in DWI images with high acceleration factors of parallel imaging. However, in Experiment 1, we relaxed the TE value to accommodate both iPAT2 and iPAT1. This will compromise the benefit of shorter TE from parallel imaging. To keep the same short TE, segmented acquisition scheme [53] can be used to collect low-noise DWI dataset without using parallel imaging. Segmented acquisition can also reduce geometric distortion, similar to parallel imaging.

Recent development of simultaneous multi-slice (SMS) techniques makes the data acquisition more efficient with only a slight SNR penalty similar to the g-factor. Experiment 2 suggested that 1D-CNN can also be used to reduce the effect of noise floor in DWI images acquired with high SMS acceleration factors. The low-noise image data can be acquired with the same DWI pulse sequence without using SMS. The SNR is higher without using SMS but only fewer slices can be acquired. Hence, it takes several scans to obtain the same amount of slices as the high-noise images using SMS.

Another possible application is for high-resolution DWI. One of the challenges for high-resolution DWI is the noise floor. However, the noise floor problem can be mitigated by constructing images from multiple thicker slices shifting in the slice-select direction [54]. The thicker slices have higher SNR. Assuming the noise distribution in the thick-slice images is Gaussian, which is a good approximation at high SNR, the noise remains Gaussian in the reconstructed image, which is a linear combination of a series of thick-slice images. Hence, the reconstructed images can be used as the ground truth for training the 1D-CNN model to correct for the noise floor in normally acquired high-resolution DWI images.

For all these applications, since the high-noise and low-noise images cannot be reconstructed from the same acquisition as in the SoS/SENSE1 case, it is critical to minimize the difference between the high-noise image and low-noise image except for the difference in SNR. An important post-processing step is to coregister the high- and low-noise datasets so that the corresponding voxels match well. However, it is not required that the high-noise images and low-noise images are acquired with the same type of sequences.

### Limitations

In the current version of 1D-CNN model, all voxel-wise time courses are used for training the model, which does not account for noise variation with respect to tissue properties and spatial locations. An improvement could be made by dividing the image into patches that have similar noise characteristics and applying 1D-CNN separately on each patch. Another limitation is the image quality of the reference itself. For instance, the SENSE1 images were used as the ground truth for training the 1D-CNN model. However, SENSE1 images also contain noise. In fact, the SNR of SENSE1 images can be very low at high b-values, at which the most severe SoS noise amplification occurs. As a result, the effect of denoising is compromised. In addition, there is a possibility of mismatch between the low-noise and high-noise dataset pair. While the SENSE1 datasets and SoS datasets were perfectly matched because they came from the same base image, it is more common in practice that low-noise and high-noise datasets are acquired separately. The effect of mismatch needs further investigation.

## 5. Conclusions

In summary, we have developed a simple 1D-CNN deep learning method to denoise DWI images and showed promising results in simulation and correcting for SoS noise in *in vivo* DWI data. By acquiring both low-noise and high-noise images of one subject, the trained model can be used to reduce noise, especially related to the rectified noise floor, in any high-noise data using the same DWI protocol and effectively convert the noise behavior in high-noise images to that of low-noise images. This method can shed light on further development of deep learning based denoising techniques for diffusion imaging.

## Ethics statement

This study involves a number of human subjects. The study was approved by the institutional ethical review board of Indiana University. Written informed consent was obtained from all participants.

## Acknowledgements

SVB was supported by an NSF SBE Postdoctoral Research Fellowship, #2004877.

